# Single-cell atlas of progressive supranuclear palsy reveals a distinct hybrid glial cell population

**DOI:** 10.1101/2021.04.11.439393

**Authors:** Abhijeet Sharma, Won-Min Song, Kurt Farrell, Kristen Whitney, Bin Zhang, John F. Crary, Ana C. Pereira

## Abstract

Progressive supranuclear palsy (PSP) is a neurodegenerative disorder whose molecular complexity at a single cell level has not been evaluated. Here we analyzed 45,559 high quality nuclei from the subthalamic nucleus and associated basal ganglia regions from post-mortem human PSP brains with varying degrees of tau pathology compared to controls (*n*=3 per group). We identified novel astrocyte-oligodendrocyte hybrid cell populations that overexpress neurotropic factors in conjunction with suppression of the unfolded protein response pathway. Notably, trajectory analysis identified subpopulations of hybrid cells with distinct astrocytic, oligodendrocytic and hybrid molecular states that change from a neuroprotective hybrid cell to an astrocytic cell with impaired homeostatic function in PSP. Our single nucleus transcriptomic data provides insights into the cell-type-specific contributions to the disease for investigating the molecular and cellular basis of PSP.

## Introduction

Progressive supranuclear palsy (PSP) is a neurodegenerative disorder, estimated to affect five in every 100,000 people, being the most common primary tauopathy^1, 2^. PSP clinically manifests with vertical supranuclear gaze palsy, loss of balance, progressive rigidity and cognitive impairment that often initiates with executive dysfunction^1, 2^. PSP is characterized neuropathologically by deposition of abnormal hyperphosphorylated tau (ptau) in neurons and glia. Diagnostic features are neuronal loss and the presence of tau-positive neurofibrillary tangles (NFTs) and tufted astrocytes (TA), and are supported by the presence of pre-tangles, neuropil threads and oligodendroglial coiled bodies predominantly in three out of four brain regions of the basal ganglia and brainstem: globus pallidus, substantia nigra, pontine nuclei and the subthalamic nucleus^3–7^. The subthalamic nucleus is one of the first, and the most severely affected brain region in this disease^8^.

PSP is largely a sporadic disease, with some reported cases of autosomal dominant mutations in the *MAPT* gene^9^. Genome wide association studies (GWAS) have identified a common haplotype in the 17q21.31 *MAPT* locus as the major genetic risk factor, however the relationship between this genetic risk and disease pathogenesis remains a critical knowledge gap^10–12^. Several PSP-associated risk alleles outside of the *MAPT* locus have been identified, including a single nucleotide polymorphism in the EIF2AK3 gene, which encodes a protein critical for the unfolded protein response (UPR)^10, 12^. UPR activation occurs in multiple neurodegenerative diseases, including Alzheimer’s disease, Parkinson’s disease, and PSP, however the role is unclear^13, 14^.

Bulk transcriptomic analyses of cortical tissue without widespread cell loss from PSP has identified transcriptional changes that potentially drive distinct cell-specific tau aggregation^15^. The study also identified a positive association of NFTs with synaptic genes and TA with microglial gene-enriched immune network^15^. This study implicates diverse molecular mechanisms underlying cell-type specific vulnerability to tau aggregation and degeneration. But, the extent to which this represents differences in the relative abundance of cells or cell specific gene expression changes cannot be ascertained with bulk transcriptomic data.

Recent studies in post-mortem AD tissue using single cell sequencing have identified cell-type specific association with Aβ and tau pathology and identification vulnerable cell populations at a single cell level has greatly improved our understanding of disease mechanisms in Alzheimer’s disease (AD)^16–18^. There is a clear need to understand cell-type-specific gene regulatory changes at a single cell level in PSP.

Here, we have characterized the transcriptional changes and cellular diversity of the subthalamic nucleus and basal ganglia structures of PSP brains using single-nucleus sequencing (DroNc-Seq). We identified novel subpopulations of astrocyte-oligodendrocyte hybrid cells marked by secretory proteins that regulate (re)myelination and synaptic function. We also used trajectory analysis to identify a potential neuroprotective hybrid cell state that might represent transitionary molecular state from oligodendrocyte-like to astrocyte-like cells.

## Results

### Single cell analysis in progressive supranuclear palsy identifies a novel astrocyte-oligodendrocyte cell-type

To investigate the cellular diversity and cell-type specific disease-related changes occurring in PSP, we isolated nuclei from fresh frozen post-mortem human brain tissue from three PSP cases and three age-matched controls (**Fig.1a**). Case status and severity was assessed using immunohistochemistry of formalin fixed paraffin embedded tissue using antisera recognizing hyperphosphorylated tau (p-tau) at Ser202/Thr305 (AT8) and confirmed the pathological status with varying burdens of neurofibrillary tangles (NFTs), tufted astrocyte and coiled bodies in the PSP cases. Staining showed distinct differences in p-tau pathology (**Supplemental Fig.1a**). Computer assisted positive pixel burden assessments demonstrated differences in burden among cases (Supplemental Fig.1b). Qualitative assessment of these cases revealed that PSP2 had a higher proportion of glial and oligodendrocyte tau pathology than a predominantly neuronal tau pathology in PSP1 and PSP3 (**Supplemental Fig.1**). We performed DroNc-Seq on the 10X platform with these cases and controls and found that two PSP samples showed preferentially higher mitochondrial rates than the controls, and retained less than 50% of all cells with the conventional < 10% threshold. As these may have arisen from the neurodegenerative changes that typify PSP, we applied an adaptive mitochondrial rate threshold to balance the number of detected genes (see methods, **Supplemental Fig.2a**). Further filtering for low depths and imputing for dropout reads resulted in 45,559 high quality nuclei profiles with a median 1,041 genes detected per nucleus.

**Figure 1.**
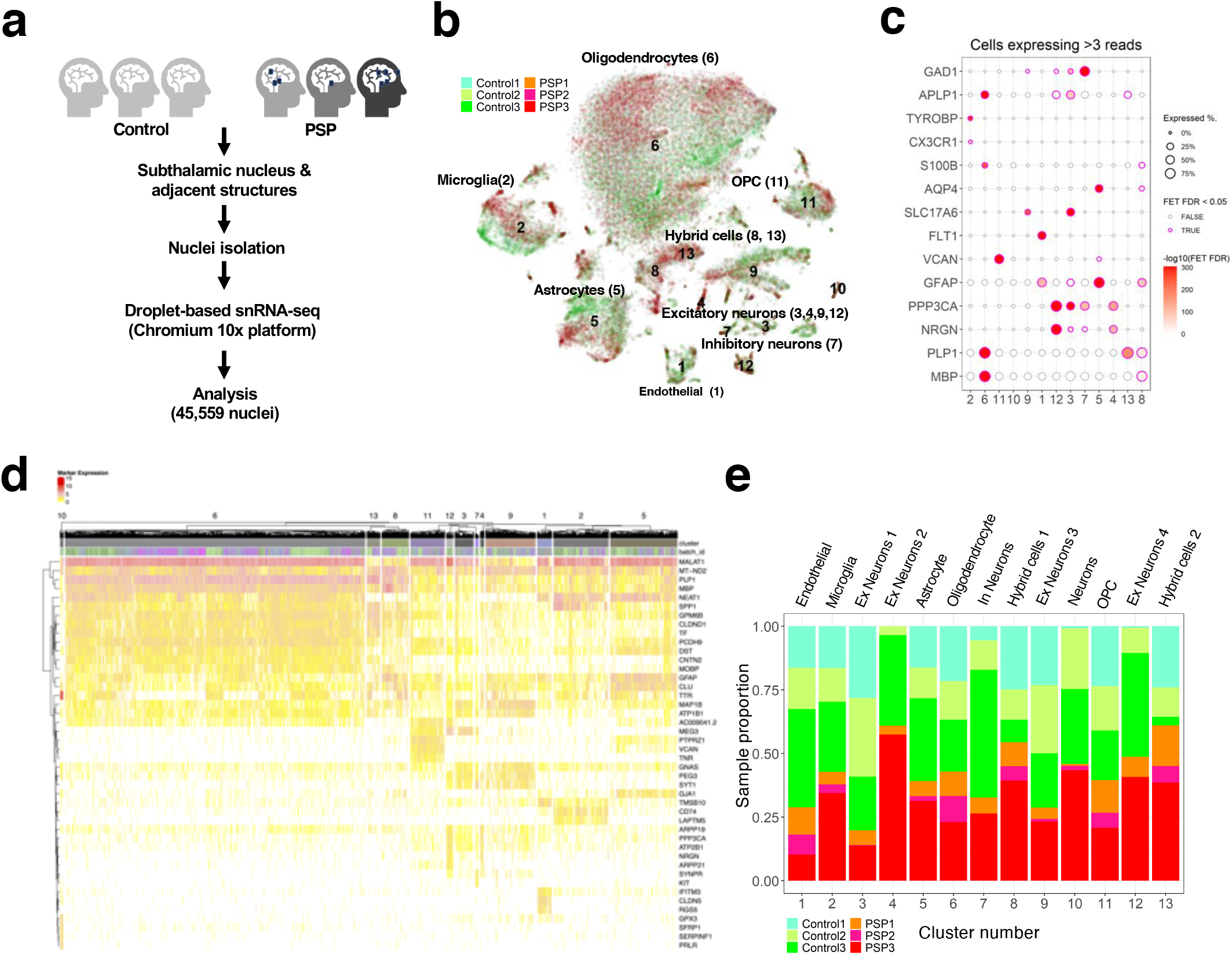
Single-nuclei sequencing of human post-mortem PSP brain identifies cell-type specific marker genes and hybrid astrocyte-oligodendrocyte cell type. **(a)** Schematic of experimental design and RNA-seq workflow. (**b)** tSNE projection of cell populations colored by individual of origin. **(c)** Marker gene expression in subclusters. A cell was deemed expressing a gene if there are more than 3 reads. The dot size depicts proportion of cells in each subcluster expressing the marker gene, and heatmap depicts enrichment of marker expressing cells in the subcluster by −log10(FET FDR). Significant enrichments are marked by magenta borders. **(e)** Hierarchical clustering of clusters and heatmap of top five marker genes in clusters. **(e)** Relative abundance of individual cell clusters.

Next, nuclei expression profiles were clustered using a two-step method based on k-nearest neighbor approach and mapped to individual cell types (**Supplemental Fig.2b, 2c)**. We identified all major cell types based on previously established cell-type-specific marker genes: excitatory neurons (NRGN, SLC17a6), inhibitory neurons (GAD1), astrocytes (AQP4), oligodendrocytes (PLP1), microglia (CX3CR1, TYROBP), oligodendrocyte progenitor cells (VCAN) and endothelial cells (FLT1) (**Fig.1b, 1c**). In addition to all major cell types, we also identified two novel hybrid astrocyte-oligodendrocyte populations (populations 8,13), that display expression of astrocytic and oligodendrocytic marker genes (**Fig. 1c, 1d**).

We compared the relative abundance of these cell types between PSP cases and controls. Analysis of relative sample-wise cell abundance in non-neuronal cell populations driven by representation from all samples revealed low relative abundance of endothelial cells in PSP (**Fig.1e**). In the neuronal populations, two excitatory neuronal populations (Ex neuron1, Ex neuron3) and the inhibitory neuronal population (In neuron) had low relative abundance in PSP (**Fig.1e**). In contrast, abundance of Ex neuron 2 and Ex neuron4 (clusters 4 and 12) were comparable in PSP and controls (**Fig.1e**). However, these differences were mostly driven by samples with the largest cell counts: Control3 and PSP3. Relative abundances of the two novel astrocyte-oligodendrocyte hybrid populations (clusters 8 and 13) were comparable to PSP with contributions from all three PSP cases (**Fig.1e**).

To better understand the molecular features that define these clusters, we identified marker genes for each subpopulation using *findMarkers* function in *scran* package. We applied FDR < 0.05 and foldchange > 1.2 to identify over-expressed genes in each cluster (**Fig.1d**). Among the neuronal clusters, Ex neuron2 and Ex neuron4 (Clusters 4 and 12) with no apparent neurodegeneration in PSP were marked by PPP3CA (PP2B), one of the major phosphatases that dephosphorylates p-tau in the brain ^19–21^ and, ARPP21 and ARPP19, genes that mediate protein phosphorylation through cAMP (**Fig.1d)**.

Next, we focused on the two novel astrocyte-oligodendrocyte hybrid populations (clusters 8 and 13) whose relative abundances were comparable in PSP with contributions from all three PSP cases (**Fig.1e**). Cluster 8 is marked by the oligodendrocyte marker genes PLP1 and MBP, and astrocytic marker GFAP (**Fig.1c, 1d**). Cluster 13 is marked by oligodendrocytic genes *PLP1, APLP1* and astrocytic gene S100B (**Fig.1c, 1d**). Gene ontology analysis of the biological function of markers of cluster 8 (t-test FDR < 0.05, |fold change (FC)| > 2) were enriched for peptide chain elongation (Fisher’s Exact Test (FET) FDR = 4.12E-10, 34.1 enrichment fold-change (EFC)) and myelin sheath (FET FDR = 7.49E-05, 17.4 EFC) and markers for cluster13 enriched for myelin sheath (FET FDR = 4.12E-10, 34.1 EFC) (**Supplemental Fig. 2e**).

### Differential gene expression analysis identifies common and unique pathways affected in PSP

We identified and compared major pathways represented by DEGs in neuronal clusters. In neuronal clusters ExNeu2 and ExNeu4 (Cluster 4 and 12), gene expression changes corresponded to several similar pathways that modulate calcium mediated signaling. Pathways are represented by reduced activity of nNOS signaling, cAMP signaling and calcium signaling (**Fig.2b, 2d)**, with significant down regulation of calmodulin (CALM1) and calmodulin-dependent protein kinase II (CAMKIIA) **(Fig.2a, 2c)**. In contrast, Ex neuron1 (cluster 3) was represented by reduced synaptogenesis signaling and dysregulation of protein ubiquitination pathways, and Ex neuron3 (cluster9) was represented by dysregulation of glutamate receptor signaling **(Fig.2f, 2h)**.

**Figure 2.**
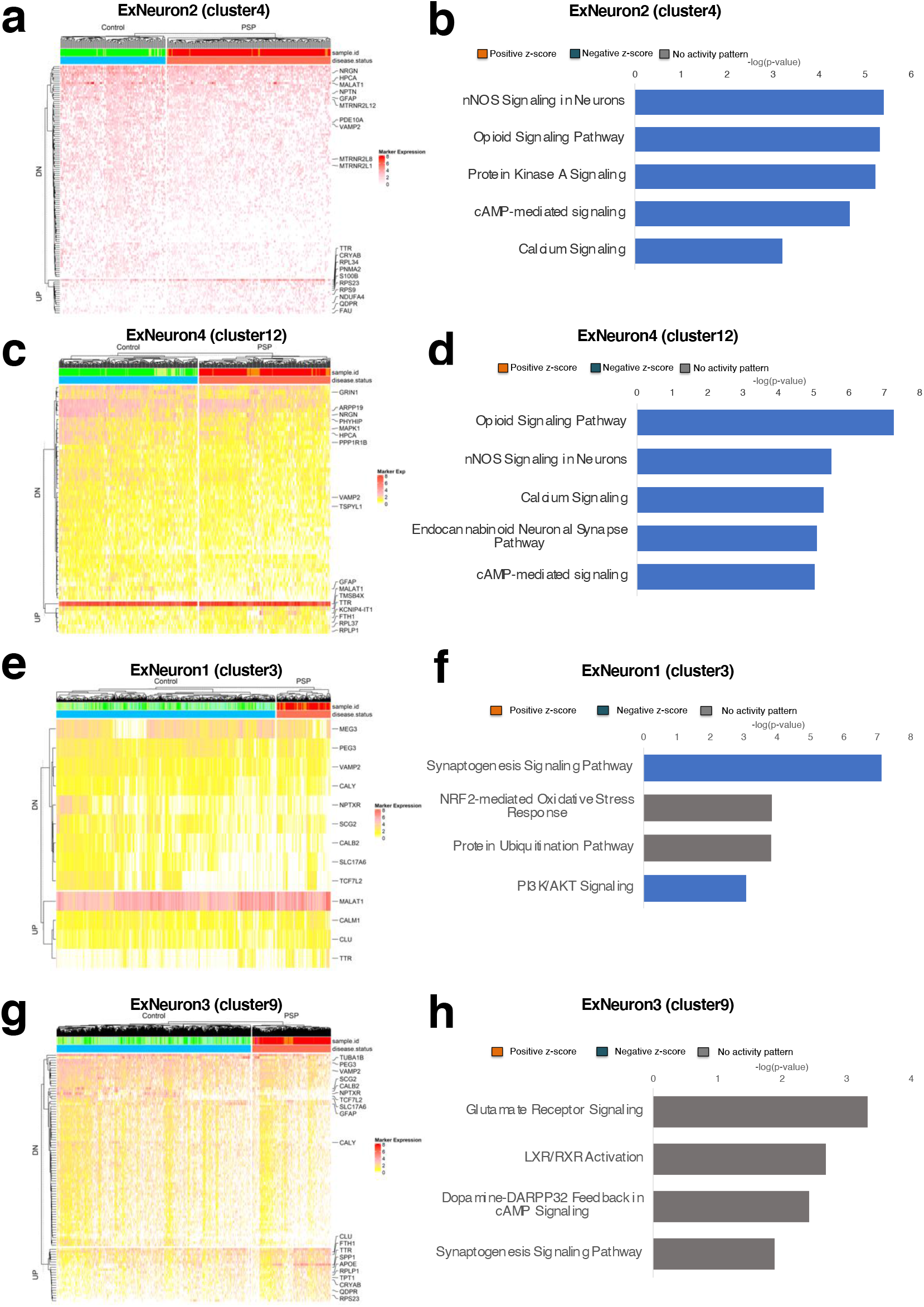
Cell-type specific pathways and differentially expressed genes (DEGs) in neuronal clusters. **(a)** Hierarchical clustering of log fold change of DEGs (log fold change >1.2 and FDR < 0.05) between PSP and control cells of neuronal cluster 4. **(b)** Canonical pathways derived from ingenuity pathway analysis (IPA) for significantly dysregulated genes in neuronal cluster (cluster 4) in PSP brains. p-values generated from Fisher’s exact test. Orange bars predict an overall increase in the activity of the pathway while blue bars indicate a prediction of an overall decrease in activity. **(c)** Hierarchical clustering of log fold change of DEGs (log fold change >1.2 and FDR < 0.05) between PSP and control cells of neuronal cluster 12. **(d)** Canonical pathways derived from ingenuity pathway analysis (IPA) for significantly dysregulated genes in neuronal cluster (cluster 12) in PSP brains. p-values generated from Fisher’s exact test. Orange bars predict an overall increase in the activity of the pathway while blue bars indicate a prediction of an overall decrease in activity. (**e)** Hierarchical clustering of log fold change of DEGs (log fold change >1.2 and FDR < 0.05) between PSP and control cells of neuronal cluster 3. (**f)** Canonical pathways derived from ingenuity pathway analysis (IPA) for significantly dysregulated genes in neuronal cluster (cluster 3) in PSP brains. p-values generated from Fisher’s exact test. Orange bars predict an overall increase in the activity of the pathway while blue bars indicate a prediction of an overall decrease in activity. **(g)** Hierarchical clustering of log fold change of DEGs (log fold change >1.2 and FDR < 0.05) between PSP and control cells of neuronal cluster 9. **(h)** Canonical pathways derived from ingenuity pathway analysis (IPA) for significantly dysregulated genes in neuronal cluster (cluster 9) in PSP brains. p-values generated from Fisher’s exact test. Orange bars predict an overall increase in the activity of the pathway while blue bars indicate a prediction of an overall decrease in activity.

In microglia (cluster 2), the top pathway from gene expression changes occurring mapped to activation of EIF2 signaling, with upregulation of a number of ribosomal genes (**Fig.3a, 3b**). Expression changes also corresponded to activation of neuroinflammatory signaling, with increased expression of B2M (**Fig.3a, 3b**), a gene associated with activated microglia (17). In endothelial cells (cluster 1), top pathways included caveolar-mediated endocytosis, EIF2 signaling and antigen presentation pathway (**Fig.3d)**.

**Figure 3.**
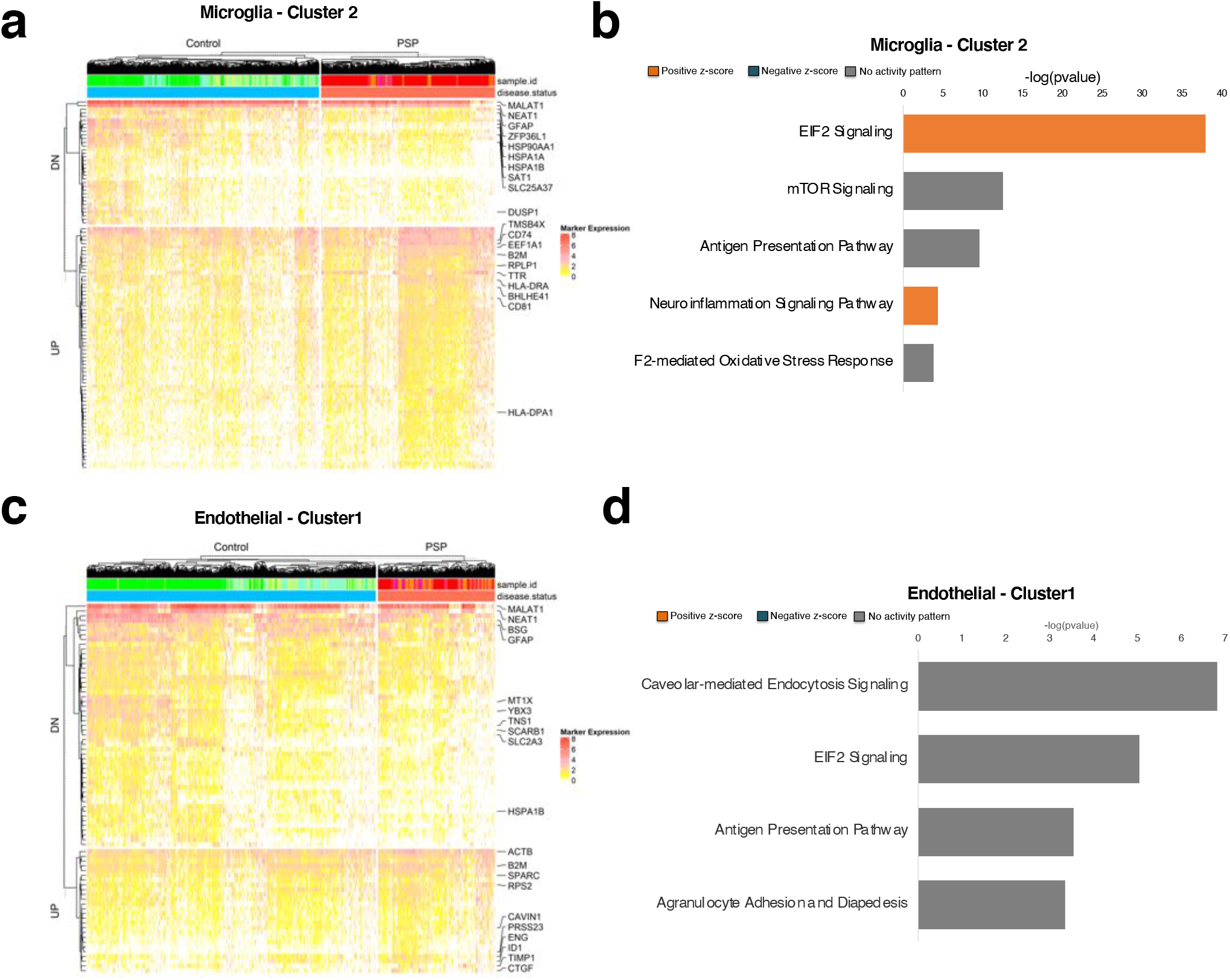
Cell-type specific pathways and DEGs in non-neuronal clusters. **(a)** Hierarchical clustering of log fold change of DEGs (log fold change >1.2 and FDR < 0.05) between PSP and control cells of microglia (cluster 2). **(b)** Canonical pathways derived from ingenuity pathway analysis (IPA) for significantly dysregulated genes in microglia (cluster 2) in PSP brains. p-values generated from Fisher’s exact test. Orange bars predict an overall increase in the activity of the pathway while blue bars indicate a prediction of an overall decrease in activity. **(c)** Hierarchical clustering of log fold change of DEGs (log fold change >1.2 and FDR < 0.05) between PSP and control cells of endothelial cells (cluster 1) **(d)** Canonical pathways derived from ingenuity pathway analysis (IPA) for significantly dysregulated genes in endothelial cells (cluster 1) in PSP brains. p-values generated from Fisher’s exact test. Orange bars predict an overall increase in the activity of the pathway while blue bars indicate a prediction of an overall decrease in activity.

We next investigated the two novel astrocyte-oligodendrocyte hybrid populations and their celltype specific gene expression changes in comparison to astrocyte and oligodendrocyte cell clusters. To validate expression of the marker genes at a single cell level, we performed immunofluorescence targeting GFAP and PLP1 and identified co-expression of these markers in hybrid cells in control and PSP tissue (**Fig. 4a**). Common enriched pathways from expression changes in astrocyte, oligodendrocyte and hybrid cell clusters in PSP are represented by pathways controlling cholesterol metabolism and (re)myelination after injury (**Fig.4c-f**). Changes in gene expression in clusters 5, 8 and 13 associated with activation of LXR/RXR signaling, with increased expression of apolipoprotein E (ApoE). Pathways from differentially expressed genes (DEGs) in astrocytes (cluster 5) are associated with dysregulation of oxidative stress, endocytosis, protein ubiquitination and downregulation of IL-6 signaling (**Fig.4c**). Pathways from DEGs in oligodendrocytes (cluster 6) are associated with dysregulation of neuregulin signaling (**Fig.4d**) and over expression of *Erbin*, a gene critical for remyelination of regenerated axons after injury ^22, 23^. Oligodendrocyte cluster 6 also show increased expression of QDPR, whose overexpression has previously been shown in an AD-pathology associated white matter oligodendrocyte subpopulation ^16^. In hybrid clusters, DEGs from hybrid2 (cluster 13) were represented by activation of cholesterol homeostasis and Bex2 signaling (**Fig.4f**). Interestingly, pathways from DEGs in hybrid1 (cluster 8) showed significant association with reduced EIF2 signaling and dysregulation of mTOR signaling (**Fig.4e**). We also identified marker genes of cluster 8 that were also overexpressed in PSP and identified several secreted proteins with known neuroprotective functions. In PSP, hybrid1 (cluster 8) cells overexpress osteopontin (SPP1), clusterin (CLU) and metalothionein2A (MT2A) (**Fig 4b**).

**Figure 4.**
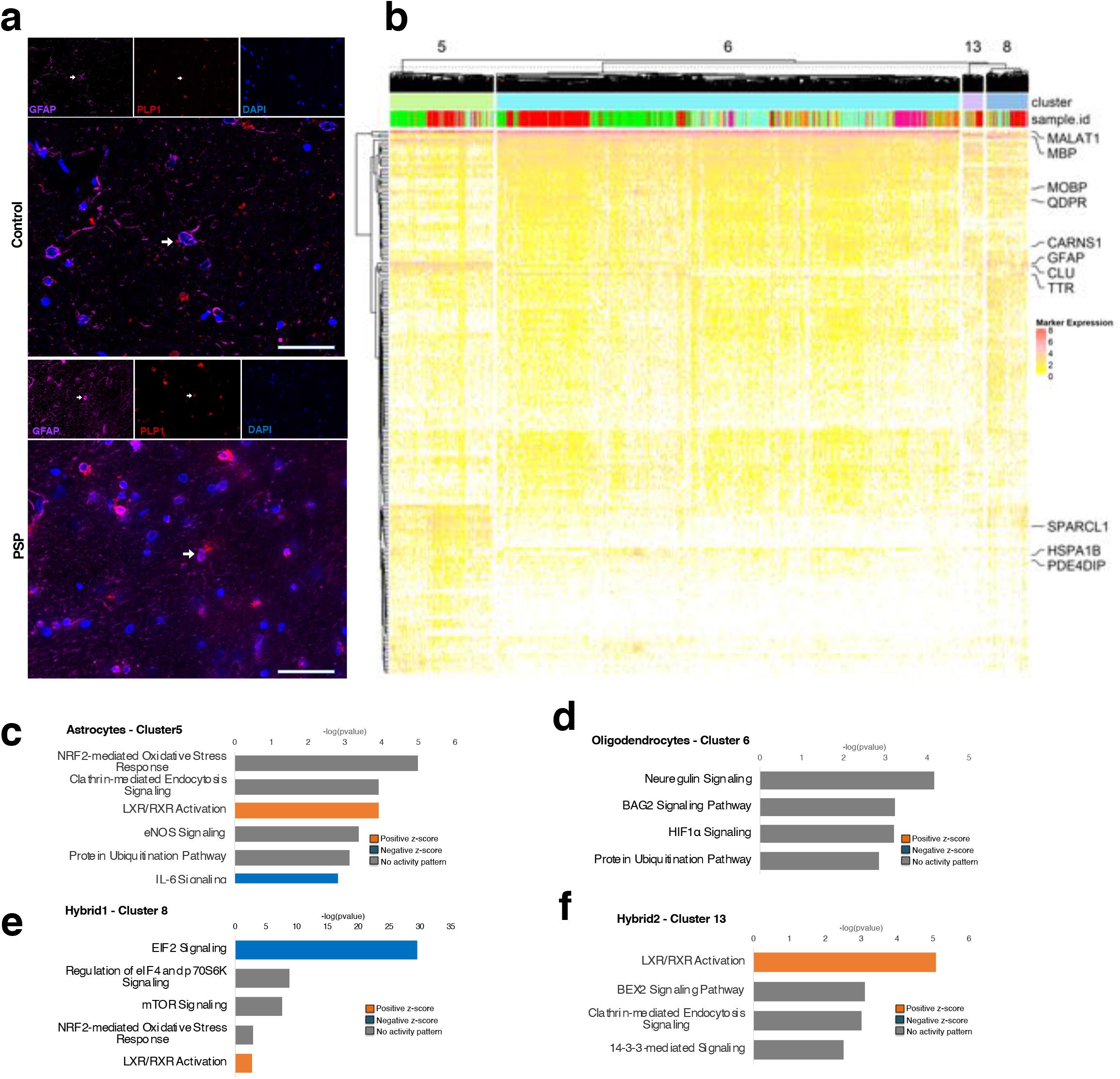
Common and unique gene expression changes in hybrid cell populations. **(a)** Representative images of hybrid cells in control and PSP sections from the subthalamic nucleus stained with GFAP antibody (magenta) and *in situ* probe for PLP1 (red). **(b)** Hierarchical clustering of log fold change of DEGs (log fold change >1.2 and FDR < 0.05) between PSP and control cells of astrocyte (cluster 5), oligodendrocyte (cluster 6) and hybrid clusters (Cluster 8 and 13). **(c)** Canonical pathways derived from ingenuity pathway analysis (IPA) for significantly dysregulated genes in astrocytes (cluster 5) in PSP brains. p-values generated from Fisher’s exact test. Orange bars predict an overall increase in the activity of the pathway while blue bars indicate a prediction of an overall decrease in activity. **(d)** Canonical pathways derived from ingenuity pathway analysis (IPA) for significantly dysregulated genes in oligodendrocytes (cluster 6) in PSP brains. p-values generated from Fisher’s exact test. Orange bars predict an overall increase in the activity of the pathway while blue bars indicate a prediction of an overall decrease in activity. **(e)** Canonical pathways derived from ingenuity pathway analysis (IPA) for significantly dysregulated genes in the hybrid cluster 8 in PSP brains. p-values generated from Fisher’s exact test. Orange bars predict an overall increase in the activity of the pathway while blue bars indicate a prediction of an overall decrease in activity. **(f)** Canonical pathways derived from ingenuity pathway analysis (IPA) for significantly dysregulated genes in the hybrid cluster 13 in PSP brains. p-values generated from Fisher’s exact test. Orange bars predict an overall increase in the activity of the pathway while blue bars indicate a prediction of an overall decrease in activity.

### Subcluster and trajectory analysis identifies an intermediate hybrid cell-state in oligodendrocyte-like to astrocyte-like molecular transitions

To further explore the diversity of the hybrid cell clusters and identify molecular changes resulting in cell-state changes, we subclustered the hybrid cell populations and performed trajectory analysis ^24^. Subcluster analysis of hybrid1 and hybrid2 (clusters 8 and 13) revealed seven clusters S1-S7 (**Fig.5a**). We then identified marker genes for the different subclusters (FDR < 0.05, |fold change| > 2 and requiring at least 25% of subcluster-pairwise comparisons to show significant differential expressions) (**Fig.5b**). Interestingly, two subclusters S2 and S6 are distinctly marked by oligodendrocyte markers PLP1 and MBP with no astrocytic marker expression (**Fig.5b**) and two subclusters S7 and S4 are marked by astrocytic markers GFAP and GJA1, with no oligodendrocytic marker expression. However, two subclusters S1 and S5 are marked by both oligodendrocytic and astrocytic genes PLP1, MBP and GFAP.

**Figure 5.**
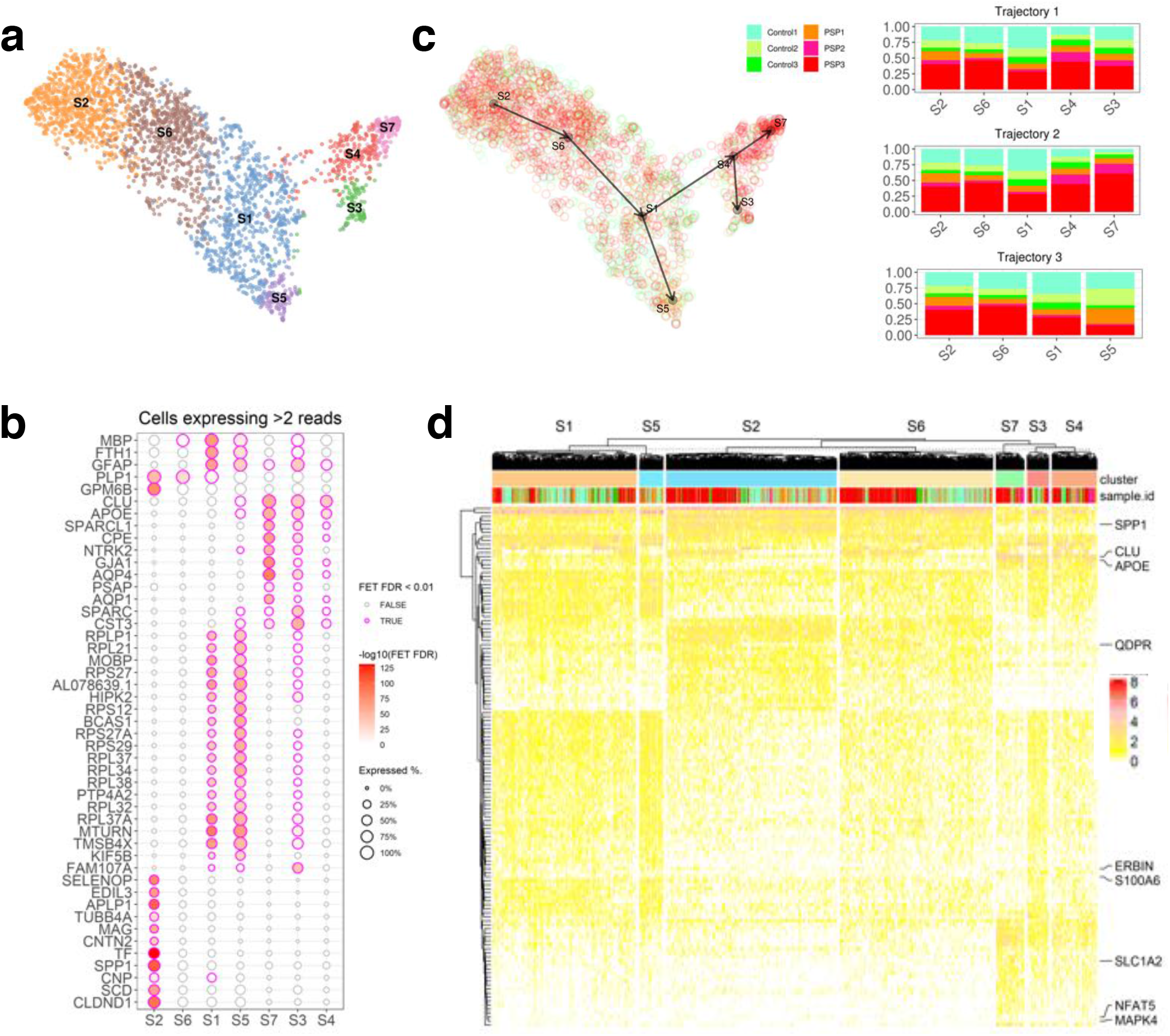
Subclustering hybrid cells reveals distinct oligodendrocyte-like, hybrid and astrocyte-like molecular states. **(a)** UMAP plot of subclusters of hybrid clusters 8 and 13. **(b)** Expressed marker genes in different subclusters. For each gene, a cell was deemed expressing a gene if there are more than 2 reads. The dot size depicts proportion of cells in each subcluster expressing the marker gene, and heatmap depicts enrichment of marker expressing cells in the subcluster by −log10(FET FDR). Significant enrichments are marked by magenta borders. **(c)** Dot and arrow plots depict inferred cell trajectories across the subclusters from S1 to S7. Cell populations are colored by individual of origin, trajectories are denoted as T1, T2 and T3, and are marked at the terminal clusters. Sample proportions of cells in subclusters are aligned by trajectories. **(d)**. Hierarchical clustering of log fold change of DEGs (log fold change >1.2 and FDR <0.05) between PSP and control cells of hybrid subclusters.

We next wanted to investigate if the hybrid subpopulation represents an intermediate molecular state between ‘oligodendrocyte-like’ and ‘astrocyte-like’ cell types. To identify potential cell-state changes, we performed trajectory analysis on the subclusters (**Fig.5c**). We applied *slingshot* (R package v1.8.0) to perform trajectory inference on the subclusters to identify pseudo-temporal dynamics among the hybrid cells. Trajectory analysis yielded three trajectories. All three trajectories indicated a change in cell-state from S2 and S6 ‘oligodendrocyte-like’ subpopulations to the hybrid S1 subpopulation (**Fig.5C**). Trajectory 1 and trajectory 2 shows an inferred trajectory from the hybrid S1 cell to ‘astrocyte-like’ S4, S7 and S3 subpopulations (**Fig.5c**) and trajectory 3 to another hybrid subpopulation S5 defined by marker genes similar to the hybrid S1 population (**Fig.5b**). Interestingly, hybrid cell populations S1 and S5 are marked by *Bcas1*, a marker of transient remyelinating oligodendrocytes (**Fig.5b**).

Sample proportions of cells in subclusters aligned by trajectories indicate an increase in relative abundance of subpopulations S7 and S4 and relative deficiency in hybrid subpopulations S1 and S5 in PSP brains driven by changes in cell numbers from all individuals (**Fig.5d**). Analysis of differentially expressed genes revealed significantly elevated expression of Qdpr, Spp1, Clu, ApoE and S100A6 in hybrid subcluster S1 (**Fig.3e**) and reduced expression of Nrxn1 and Slc1a2, genes regulating homeostatic astrocyte function in astrocyte-like subpopulation S7 (**Fig.5e**).

## Discussion

Tau pathology is a common feature of a number of neurodegenerative disorders. While AD is the most common, PSP is the most common primary amyloid-independent tauopathy. Understanding the cellular and molecular alteration in PSP has the potential to not only pinpoint disease-specific pathways but also identify common pathogenic mechanisms of tau-mediated neurodegeneration. The data from this study highlight the distinct changes occurring in all the major cell types in brain regions vulnerable to PSP. In this study, we have uncovered an astrocyte-oligodendrocyte hybrid cell population with potential neuroprotective properties whose numbers are reduced in PSP. Further, these data also shows distinct neuronal populations, some of which have an adaptive response against neurodegeneration.

Excitatory neuronal loss is a pervasive phenotype in several neurodegenerative disorders. We have identified two excitatory neuronal populations that do not have reduced abundances in PSP. Both excitatory neuronal populations (Ex neuron2 and Ex neuron4) are marked by PP2B, ARPP19 and ARPP21. PP2B (also known as calcineurin) is the only phosphatase directly modulated by calcium to control synaptic transmission and other neuronal processes^25^. PP2B is one of the major phosphatases that dephosphorylates p-tau in the brain^19–21^. Expression and activity of PP2B is reduced in brains of patients with Alzheimer disease^26^ and reduced PP2B activity is directly associated with increased p-tau in chronic traumatic encephalopathy (CTE)^27^. ARPP19 and ARPP21 are substrates for calcium dependent cAMP-dependent protein kinase (PKA) and expression of ARPP19 is reduced in AD brains^28^.

Differential gene expression analysis indicated that these clusters (Ex neuron2 and Ex neuron4) have downregulation of calcium signaling and cAMP signaling pathways with significant down regulation of CALM1 and CAMKIIA. Disruption of calcium homeostasis and activation of calmodulin dependent CAMKIIA has been shown to impair synaptic morphology and trigger neuronal apoptosis in AD^29, 30^. CAMKIIA also associates with, and phosphorylates tau at multiple sites to induce a conformational change to generate paired helical fragment conformation^31–35^. The expression changes indicate an adaptive response in these excitatory neuronal populations to prevent synaptic loss and degeneration. These cell clusters are mainly comprised of cells from samples with the largest cell counts. A limitation of our study is the heterogeneity of PSP pathology in the three individuals. Whether there are any significant associations of particular neuronal subpopulations with tau pathology requires systematic stratification of the PSP variants and further analysis with a larger cohort of individuals with similar PSP tau pathology.

Single cell studies in other neurodegenerative disorders like AD and Huntington’s disease have identified hybrid cells which may represent intermediate cellular states^18, 36, 37^. However, the functional roles of these intermediate cellular states have not been elucidated. Injury and stress in the brain potentially drive molecular changes in certain cell populations to mitigate neuronal and glial damage. Astrocyte, oligodendrocyte and hybrid cell clusters in PSP brains have expression changes that are represented by pathways controlling cholesterol metabolism and (re)myelination after injury. These changes indicate that an injury response is activated in these cell types.

Genetic and pathological studies have implicated the PERK/eIF2 pathway and unfolded protein response (UPR) activation in the pathogenesis of PSP^10, 38^. Dysregulated UPR signaling driven by altered expression of ribosomal proteins can be seen in multiple cell types in PSP. Although the relationship between the unfolded protein response and autophagy has been well studied in neurons, its role in regulating cell autonomous and non-cell autonomous functions in other cell types has not been well studied. It is unclear if dysregulation of UPR is a common stress mechanism that affects multiple cell types in the brain during neurodegeneration or a response specific to PSP.

Differentially expressed genes in hybrid cells significantly map to downregulation of EIF2 signaling. Activation of the PERK/eIF2 mediated UPR signaling in astrocytes of mice infected with prion protein has been shown to drive non-cell autonomous neurodegeneration^39^. PERK activation is seen in brain regions highly affected by tau, but tau and pPERK do not overlap at a single cell level^38^. It remains unclear if accumulation of tau in astrocytes induces PERK activation. The expression changes seen in the hybrid cells in PSP suggest that they represent a non-cell autonomous stress response to neurons undergoing demyelination and degeneration. Given that the UPR pathway is the main pathway maintaining cellular homeostasis under conditions of protein aggregation and misfolding, this cell state might also represent a cell intrinsic protective mechanism in the hybrid cells to prevent tau aggregation.

The astrocyte-oligodendrocyte hybrid cells in PSP brains are marked by, and overexpress several genes that produce secretory proteins with known neuroprotective roles. SPP1 has been shown to promote repair processes after injuries in the CNS by modulating inflammatory responses^40–43^. Elevated clusterin levels has been reported in AD-vulnerable brain regions and an AD-protective variant of CLU is associated with higher expression^44,45^. Clusterin has been shown to be associated with tau in AD and primary tauopathies^46,47^ and preferentially expressed by AD pathology-associated astrocyte subpopulation^16^. Additionally, loss of CLU has been associated with exacerbated tau pathology in a mouse model of tauopathy^47^. Increased expression of metallothionein (MT) is associated with several neuronal injuries including ischemia, infection, Huntington’s disease and AD^47^. AD pathology-associated astrocytes also preferentially express several MT genes^16^. Increased expression of MT genes has also been shown in astrocyte subpopulations suggested to be neuroprotective and astroprotective in Huntington’s disease^36^. Mice deficient in MT1/2 show reduced neuronal survival after ischemic injury, impaired wound healing and astrogliosis after freeze-injury^48,49^. These gene expression changes occurring in the hybrid cells indicate that the homeostatic function of these cells is to support (re)myelination and mitigate neurodegeneration by tropic factor secretion.

Sub-clustering analysis of the hybrid cell populations revealed three distinct molecular states - a oligodendrocyte-like, astrocyte-like and a hybrid state with markers of both astrocytes and oligodendrocytes. The hybrid subpopulations are also marked by *Bcas1*. Breast-carcinoma amplified sequence 1 (Bcas1) has been shown to specifically mark a distinct population of oligodendrocytes that are present transiently during the active phase of oligodendrocyte generation and myelination in human and rodent brains^50, 51^. Bcas1^+^ oligodendrocytes have also been associated with partially remyelinating lesions in multiple sclerosis^50^. There are currently no markers for active myelination, but expression of Bcas1 as a marker of the hybrid cell subpopulations suggests a similar role for these cells in PSP brains. The spatial localization of these hybrid cells, association with tau pathology and functional role are intriguing and will be the subject of future work.

The hybrid subpopulations also overexpress S100A6 (calcylin) in PSP. S100A6 marks astrocytic progenitors in the subgranular zone^52^ and is over expressed by reactive astrocytes surrounding β amyloid plaques in AD and neurodegenerative lesions in amyotrophic lateral sclerosis (ALS)^53–55^. S100A6 can bind to Zn^2+^ to prevent Zn^2+^ induced toxicity^37, 56^. Zinc can also bind tau directly to contribute to tau toxicity independent of tau hyperphosphorylation^57, 58^. A recent report has revealed that S100A6 can also enhance the phosphatase activity of PPP5C towards p-tau^59^. This data further strengthens the idea that the hybrid subpopulation is associated with potential neuroprotective functions.

Trajectory analysis of the hybrid populations suggests a shift in relative abundance of hybrid S1 and S5 subpopulations to astrocyte-like S7 subpopulations in PSP. Changes in abundance indicate a shift from a neuroprotective hybrid cell state to a dysfunctional astrocyte-like state in PSP. However, without evidence for functional differences in the different subpopulations, distinguishing plasticity from heterogeneity is difficult. Identifying phenotypic differences between the cell states is necessary for a deeper understanding of the role of these hybrid cells in disease progression and therapy in PSP.

In conclusion, our study contributes a pioneering characterization of cell-type specific changes in PSP using snRNA seq. These results also highlight the need for a detailed functional characterization of multiple cell types to develop better models and therapeutic modalities for PSP. Our study will inform future studies to identify the mechanisms of tau accumulation, degeneration and repair in PSP.

## Acknowledgements

This work was supported by NIH R01AG063819 (ACP), R01AG064020 (ACP), Paul B. Beeson Emerging Leaders Career Development Award in Aging K76 AG054772 (ACP), R01AG054008 (JFC), R01NS095252 (JFC), NIA career development award (P50AG005138), <u>Karen Strauss Cook Research Scholar Award (ACP/JFC)</u>, the Alzheimer’s Association, Alexander Saint-Amand Scholar Award (JFC), Tau Consortium/Rainwater Foundation (JFC).

## Author contributions

A.S, A.C.P and J.F.C conceptualized and designed the experiments. A.S generated the snRNA-Seq data with support from K.F. W.S and A.S analyzed snRNA-Seq data and visualized results. K.W and K.F contributed to neuropathological data generation B.Z provided data analysis tools. All authors read and approved the final version of the manuscript.

---

**Supplemental Figure1.**
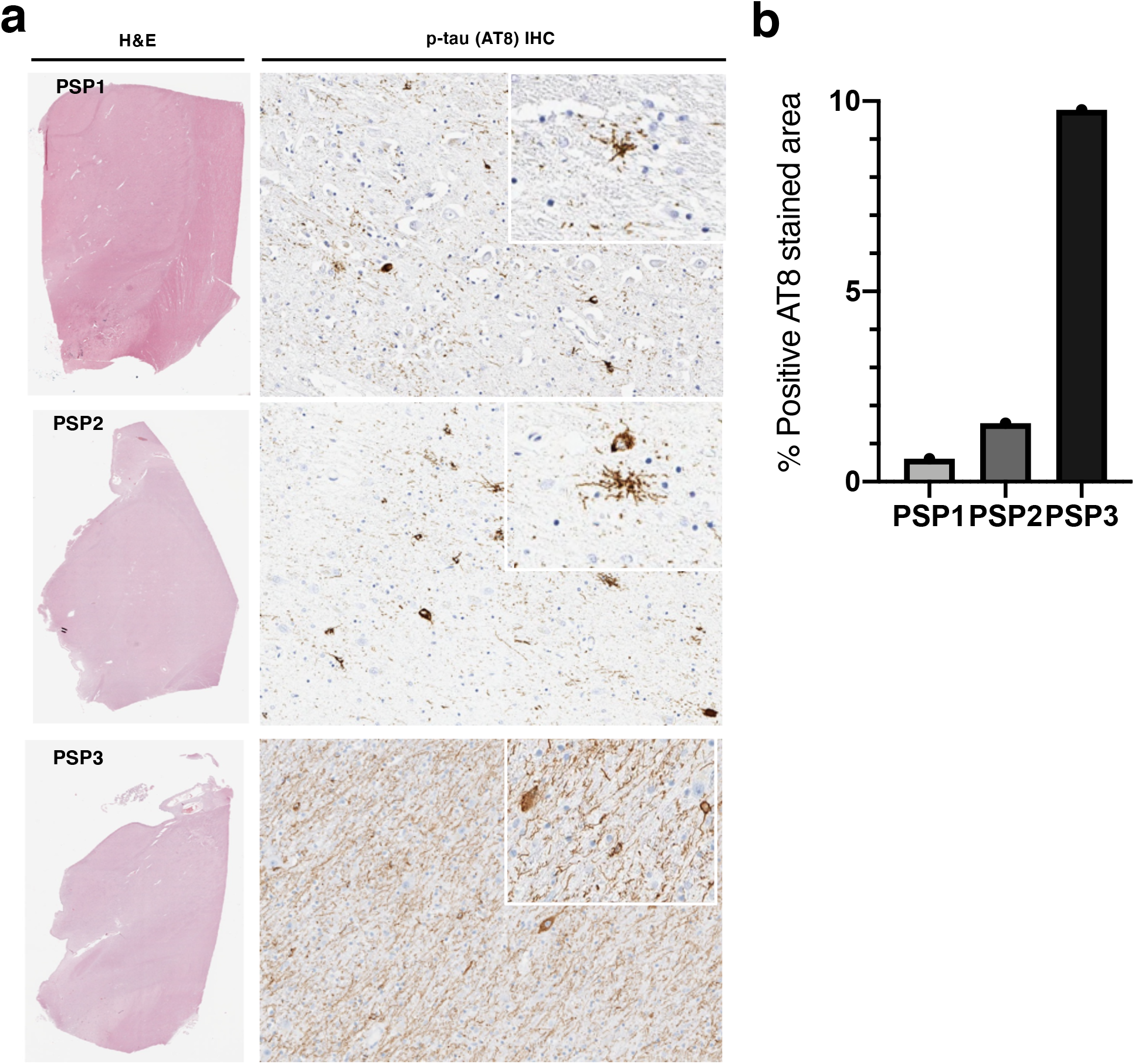
Distinct tau pathology in PSP cases. **(a) H&E** sections prepared from the contralateral hemisphere were screened for gross morphological characterization and immunostained for p-tau using AT8 antibody for characterization of tau pathology. **(b)** Quantitative pixel-level assessments of AT8 positivity as a measure of tau burden in the PSP cases.

**Supplemental Figure2.**
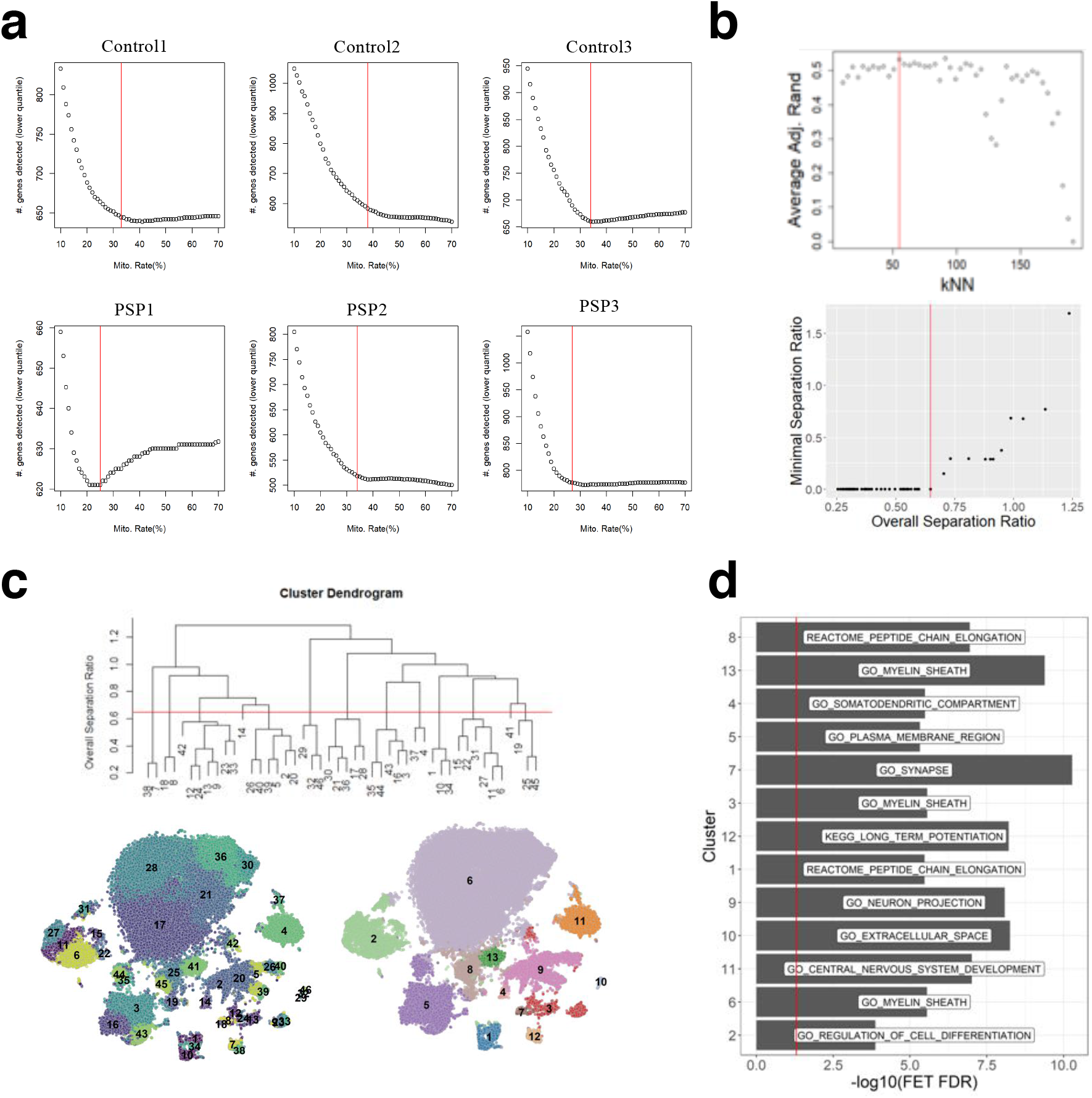
Adaptive mitochondrial thresholds per sample and identification of distinct cell clusters. **(a)**. Horizontal red lines mark the elbows as the final mitochondrial rate thresholds per sample. **(b)** Average adjusted Rand index of each clustering solution at different kNN parameters, compared to solutions at other kNN values. The red line remarks the optimal kNN value whose clustering solution shows the best overall adjusted Rand score. tSNE plot of cells from MNN-corrected data. The clusters from the first-step of kNN clustering are labeled. **(c)** Dendrogram showing similarity between the initial cell clusters, as evaluated by separation ratio. (**d)** Plot of overall separation ration (the average separation between all cell clusters merged in B), against the minimal separation ratio within clusters merged in C. The red line remarks the elbow point applied in C. tSNE plot of cells and cell clusters, refined by the separation ratio metrics in C. tSNE plot of cells and cell clusters, refined by the separation ratio metrics in C. **(e)** Top enriched functions and pathways in the cluster marker genes.

## Methods

### Dissection of Subthalamic nucleus and associated basal ganglia from frozen tissue

Postmortem specimens frozen during autopsy from control and PSP cases were obtained from the Mount Sinai Brain Bank. 3 cases of control and 3 cases of PSP were selected for snRNAseq. Tissue blocks were mounted on a cryostat and 30μm scrolls of the frozen tissue were cut to ensure equal representation of the entire block of frozen tissue. Nuceli were isolated from the scrolls immediately. A table of the cases and controls used is provided in the table below

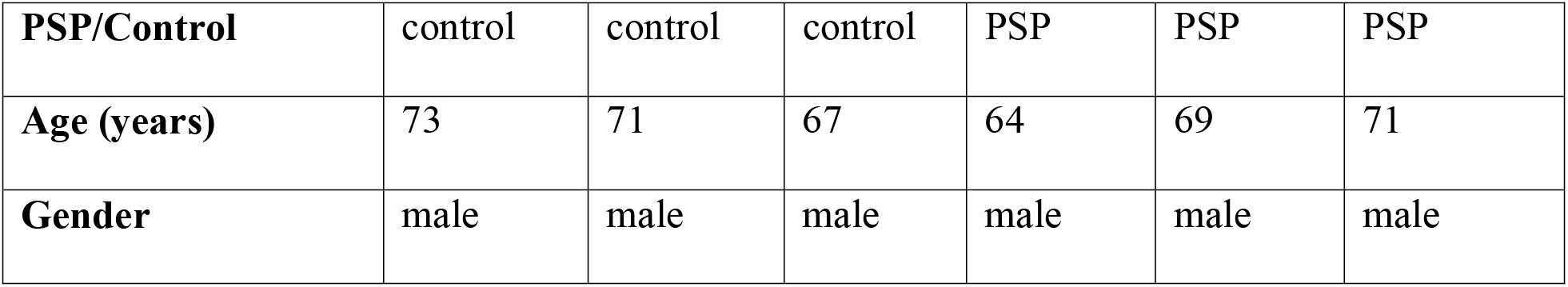

### Nuclei isolation

Tissue scrolls were homogenized in a dounce homogenizer with Nuclei EZ lysis solution. The suspension fromt each sample was filtered through a 40um Flowmi™ strainer, washed, re-filtered and resuspended in 0.05% BSA PBS containing RNAsin RNAse inhibitor. Sequencing was performed by Genewiz Inc. Nuclear suspension was counted and processed by the Chromium Controller (10x Genomics) using single Cell 3’ Reagent Kit v2 (Chromium Single Cell 3’ Library & Gel Bead Kit v2, catalog number: 120237; Chromium Single Cell A Chip Kit, 48 runs, catalog number: 120236; 10x Genomics).

### Histology, *in situ* hybridizations, and immunohistochemistry

Formalin-fixed paraffin-embedded (FFPE) tissue sections were used for H&E staining, immunohistochemistry (IHC) and *in situ* hybridizations. FFPE sections (4-5 μm) were deparaffinized, hydrated through an alcohol series and boiled in CC1 (citric acid buffer, Roche Diagnostics, Basel Switzerland) for 1 hour (pH 6) for antigen retrieval. IHC for phospo tau was performed using AT8 (pSer202/pThr205) antibody (Mouse monoclonal, 1:2000, Thermo Fisher MN1020) and visualized using the Ultraview detection kit (Roche Diagbostics) on a Benchmark XT autostainer (Ventana) according to manufactuer’s directions. Slides were counterstained with hematoxylin. For unbiased digital quantitative assessment, slides were imaged using an Aperio CS2 (Leica Biosystems, Wetzlar Germany) digital slide scanner at the Department of Pathology at Mount Sinai. Tissue sections were manually segmented on whole slide image sections, batch-processed and analyzed in QuPath (version 0.2.3, https://QuPath.github.io/). A custom pixel classifier was used to determine positive pixel count based on thresholded value for DAB intensity. Percent (%) positive pixels was automatically generated and normalized by the number of positive pixel counts to total area.

*In situ* hybridization was done using RNAscope™ multiplex fluorescent v3 (ACDbio) as per the manufacturer’s protocol in 10μm paraffin embedded, formalin-fixed tissue sections. Staining was performed with a probe to detect PLP1 expression(Cat no. 564571-C2) and GFAP antibody (rabbit polyclonal, 1:1000, Invitrogen Cat no.10019). Secondary antibody conjugated to flurophores anti-rabbit Alexa Fluor 488 and 647. Images were taken on Zeiss Axioplan with ApoTome and Aperio LSM™ slide scanner

### scRNA-seq quality control (QC) per sample

Single-cell RNA-seq (scRNA-seq) data will be QCed (scQC) by employing *scran*^60^ package (v 1.14.6). Low-quality cells were removed by outlier detection for low library size and proportion of mapped endogeneous reads with Median Absolute Deviation (MAD) > 3.

Another metric to filter out low quality cells is mitochondrial read rate (T_mito_), as a proportion of mitochondrial transcripts in each cell’s library. As PSP is a neurodegenerative disease with greater proportion of apoptotic cells than healthy control samples, we sought to apply adaptive mitochondrial rate thresholds that remove highly apoptotic cells while minimizing the loss on functional transcripts. Thus, we analyzed the number of expressed genes in cells (as a proxy for functional information) across a range of mitochondrial rate thresholds (T_mito_) between 10-70%. Per threshold, the representative number of expressed genes (N_eg_) was obtained as the lower quantile value among the filtered cells with the threshold. As the lower threshold enriches for cells with better quality, we expect that the overall number of expressed genes abruptly increases. On the other hand, the higher threshold will enforce inclusion of cells with lower number of expressed genes, thus N_eg_ will gradually reach its plateau towards the N_eg_ of all cells without any mitochondrial rate thresholding. To identify the balance between these two extremities, we looked for the elbow in the T_mito_ -vs- N_eg_ curve and obtained the adaptive threholds (see Error! Reference source not found.). The overall numbers of QCed cells show less proportion of cells from PSP remain than those from the control samples.

Per sample, doublet cells were evaluated by DoubletFinder()^61^ (v 2.0.3) from *Seurat* and doubletCells() from *scran*. Firstly, cell clusters enriched with high confidence calls from DoubletFinder (FET FDR < 0.05), and high doublet scores from doubetCells (T-test FDR < 0.05) were marked as doublet clusters and removed. Further, individual cells called with high-confidence doublets by DoubletFinder were removed. Then, the dropout reads were estimated using SAVER^62^.

### Data Merging and Batch Correction

The QCed data from individual samples were combined and corrected for batch effects by Mutual Nearest Neighbor (MNN) approach implemented in *batchelor* R package (v1.2.4) while preserving biological contents^63^. Specifically, the cells were down-scaled to match the least-sequenced batch’s coverage using “multiBatchNorm()” in batchelor package, to mitigate the differences in variances between batches^64^. We used ‘modelGeneVar()’ from the scran R package (v1.16.0) to model technical and biological variances in individual gene expressions within each batch. The technical variations were inferred by fitting variance-mean curve as the technical variation, and the deviation from the fitted variance-mean curve was modeled as biological variations with ‘modelGeneVar()’ from the scran R package (v1.16.0). These variances were combined across the batches with ‘combineVar()’ function from the scran R package (v1.16.0), and extracted variable genes exhibiting any biological variance. The MNN-based batch correction was then executed by “fastMNN()” function from batchelor package on the chosen variable genes set.

### 2-Step Unsupervised Cell Clustering and Marker Analysis

In the first step of cell clustering, we performed k-Nearest-Neighbor (kNN) graph based clustering within batch corrected feature space identified MNN approach implemented in *batchelor* package. To obtain the optimal kNN parameter to construct kNN graph, we performed walktrap clustering on each kNN □ [log(N_c_),√N_c_] where N_c_ = number of cells. We used ‘cluster_walktrap()’^65^ function implemented in *igraph* package (v 1.2.5). The clustering solutions across different kNN were then compared to each other by adjusted Rand index to evaluate degree of similarity between them. kNN=55 showed the highest overall similarity to other solution (i.e. the centroid), and was chosen as the first-step solution with 44 cell clusters.

While the first-step solution provides fine-resolutions to the cell population architecture, we sought to identify coarse-grained clusters reflecting more robust cell populations. Thus, in the second step, we evaluated inter-cluster similarity to merge some of similar cell clusters. As the distant metric between the clusters, we utilized correlation distance d_ij_ = √(2(1-ρ_ij_), where ρ_ij_ is Pearson correlation between two cells, i and j, within MNN-corrected feature space, as the cell-cell distance. Then, inter-cluster distance between clusters a and b, D_ab_, was calculated by D_ab_ = d[a,b]/[r(a) + r(b)], where d[a,b] = distance between centroid cells in clusters a and b, r(a), r(b) = radius of cluster a or b. r(a) was defined as the distance between the centroid cell of cluster a, and 90% quantile distance to other cells in the same cluster. Overall, D_ab_ represents a separation ratio between two cell clusters, where D_ab_ >1 if the centroids are farther apart than the radii of the clusters, D_ab_ < 1 otherwise. Based on the separation metric, we performed hierarchical clustering among the clusters with average linkage method, and evaluated the cutoff by inspecting the overall separation ratio (i.e. average of pairwise D_ab_) against the minimum separation ratio. We chose the elbow point in, which corresponds to average D_ab_= 0.6478. This yielded 13 cell clusters.

We performed pairwise differential expression analysis among the 13 cell clusters by utilizing *“findMarkers()”* function in *scran* package. findMarkers() first performs student t-test for all pairs of cell clusters with normalized, log-transformed expressions, then summarizes the overall p-values and FDR by averaging over the log-transformed p-values, followed by Benjamini-Hochberg correction. We applied FDR < 0.05 and fold-change > 1.2 to identify over-expressed genes in each cluster.

### Subclustering and Trajectory Inference

Two adjacent cell clusters 8 and 13 expressed markers of oligodendrocyte (*PLP1*) and astrocyte (*AQP4*) simultaneously, hence were identified as hybrid cells (denoted as H8 and H13). To further dissect transcriptome architecture of hybrid cells, we performed subclustering in these cells. The gene expressions were corrected by Mutual Nearest Neighbor (MNN) approach as described previously in ‘Data Merging and Batch Correction’ to adjust for non-biological sample effects. The adjusted data were subject to kNN-based walktrap clustering with kNN=log(N_c_) where N_c_=number of hybrid cells. This yielded 7 subclusters with distinct sample compositions. The cell markers were identified by *findMarkers()* with FDR < 0.05, |fold change| > 2 and requiring at least 25% of subcluster-pairwise comparisons to show significant differential expressions.

Then, we applied *slingshot* (R package v1.8.0) to perform trajectory inference on the subclusters to identify potential pseudo-temporal dynamics among the hybrid cells. We utilized uniform manifold approximation and projection (UMAP) in the MNN-adjusted data as the dimension reduction, onto which cell trajectories were inferred. The analysis yielded 3 trajectories.

**Table 1.** Sample information

| Category | Control1 | Control2 | Control3 | PSP1 | PSP2 | PSP3 |
| --- | --- | --- | --- | --- | --- | --- |
| **Genes before QC** | 33538 | 33538 | 33538 | 33538 | 33538 | 33538 |
| **Cells before QC** | 10907 | 8516 | 11931 | 6004 | 4221 | 13699 |
| **Genes after QC** | 20230 | 20181 | 20526 | 18543 | 16762 | 21696 |
| **Cells after QC** | 9310 | 7259 | 10266 | 3829 | 3070 | 11828 |
| **mito. rate threshold (%)** | 33 | 34 | 25 | 38 | 34 | 27 |
| **Median #. Genes** | 979 | 1056 | 898 | 1020 | 823 | 1338 |
| **QC pass rates (%)** | **85.36** | **85.24** | **86.04** | **63.77** | **72.73** | **86.34** |

## Notes

### Competing Interest Statement

The authors have declared no competing interest.

